# Paraquat Inhalation Produces Male-Biased Dopaminergic Neuron Loss and Motor Deficits in Mice: Relevance to Parkinson’s Disease

**DOI:** 10.64898/2026.11.02.761058

**Authors:** Pablo Reina-González, Jacob Boos, Abdulla Abu-Salah, Aiesha Anchan, Rachel Strazdins, Muhammet Ay, Ahmed Shah, David Chalupa, Jason Richardson, Souvarish Sarkar, Deborah A. Cory-Slechta

**Author notes:** For correspondence, please contact.

## Abstract

Long-term exposure to paraquat (PQ), a widely used herbicide, has been linked to increased risk of Parkinson’s Disease (PD) in agricultural workers and rural communities in the United States (US), findings further supported by experimental animal studies. Although banned in many countries across the globe, PQ remains widely used in the US. Inhalation of PQ has been known to induce systemic organ failure at acute concentrations. However, prolonged low-dose inhalation from agricultural spraying and pesticide drift has not been readily evaluated for its potential to contribute to neurodegenerative diseases *in vivo*. Instead, many experimental studies have relied on systemic routes of exposure, which do not effectively replicate the most common route of exposure to PQ in the environment. In the current study, Parkinsonian features were examined in mice that had been exposed via whole-body inhalation to PQ aerosol for either 40 or 60 days. Levels of PQ in brain were highest in olfactory bulb, followed by frontal cortex and then brainstem in both sexes. After 60 days of exposure, male-specific impairments in locomotor activity were observed, in addition to reductions in striatal dopamine, in substantia nigra tyrosine hydroxylase (TH) expression and number of TH+ dopaminergic neurons, the latter being considered the primary basis of PD, along with increased olfactory bulb α-synuclein levels. As such, these studies confirm the dopaminergic neurotoxicity of inhaled PQ. These findings raise significant concerns as to the contribution of PQ exposures to the continued increase in the incidence of PD in the US.

**One-sentence summary:** Paraquat inhalation in mice induces male-biased dopaminergic neuron loss and motor function deficits coupled with increased levels of α-synuclein.

## INTRODUCTION

Parkinson’s Disease (PD) is the second most prevalent age-related neurodegenerative disorder in the world after Alzheimer’s disease (AD), and notably the most rapidly increasing neurological disease [1]. Currently estimated to affect 10 million people worldwide, the global burden of PD doubled between 1990 and 2017, with some studies projecting that cases will double again by 2030 [2]. Such findings underscore the critical need to understand the mechanisms underlying PD and the environmental risk factors associated with it. The main risk factor for PD is age, with risk increasing from 1% in people over 60 to 5% in those over 85 years of age [3, 4], and given that the number of Americans over 65 is continuously escalating, a drastic rise in PD cases over the next 20 years inevitable [5, 6]. Therefore, developing effective therapeutics and public health interventions to mitigate its rise remains a priority.

PD is considered to arise from a combination of genetic and environmental factors, with environmental factors predominating [7]. One environmental factor repeatedly and strongly linked both in epidemiological studies and in animal models to increased PD risk is paraquat (PQ) [8–10], a defoliant and broad-spectrum herbicide used to clear green plant matter from soy and cotton plants during harvest season, facilitating the harvesting of grains and fiber [11, 12]. While currently banned in 70 countries, including the European Union and China, its use in the US more than doubled between 2010 and 2018, the latest year on record.

While some studies have disputed this association [13], recent work in the California Central Valley identified clear and compelling associations between working near agricultural sites utilizing PQ as well as in residential proximity to these sites, with an increased risk of PD, irrespective of sex (odds ratio = 2.08-2.15 and 1.72-1.91, respectively) while controlling for other potential exposures [14]. A 2019 EPA Human Health Risk Assessment identified potential health risks from PQ in mixers, loaders, applicators, and bystanders at the edge of the field due to drift [15]. As a result, PQ use is intended to be restricted to certified applicators wearing appropriate personal protective equipment with restricted re-entry intervals at application sites. Despite these risk mitigation measures, as the California Central Valley study suggests, people living in rural areas near application sites remain at a greater risk of PD than the rest of the population, indicating that inhalation via pesticide drift remains a relevant route of exposure [14]. This is particularly salient given recent data reported by the EPA indicating that the aerosolization of PQ is dramatically higher than initially believed (https://www.epa.gov/pesticides/epa-updates-review-potential-paraquat-volatilization-and-plans-request-additional-data) and would contribute to both inhalation and dermal exposures.

Furthermore, studies in animal models have repeatedly reported dopaminergic neurotoxicity in response to PQ exposure, including loss of dopaminergic neurons in the substantia nigra pars compacta (SNc), an effect considered the ‘gold standard’ pathological feature of PD. However, these studies have utilized routes of exposure other than inhalation, such as dietary intake, drinking water, or injection, prompting criticism that consequent effects do not accurately reflect the conditions under which human exposure to PQ is most likely to occur.

We previously implemented a short-term inhalational PQ exposure paradigm in adult mice to assess the potential brain uptake [16]. It established the highest levels in the olfactory bulb, consistent with nose-to-brain translocation of PQ. It was also detected in other brain regions, including the striatum, midbrain, cerebellum, and brainstem. In addition, male-specific impairments were found in an olfactory discrimination paradigm conducted 135 days post-exposure, demonstrating PQ’s capacity for nose-to-brain transport, i.e., to circumvent the blood-brain barrier (BBB) through the nasal epithelium, leading to olfactory deficits such as hyposmia or anosmia in male mice [17]. Olfactory dysfunction is considered a component of the prodromal stage of PD [18].

However, as noted, the primary feature of PD is the loss of dopaminergic neurons in the SNc, an effect thought to specifically underlie the movement disorders characterizing this disease [19, 20]. The current study sought to extend this inhalationmodel by utilizing longer-duration PQ inhalational exposures and assessing multiple characteristics of the PD phenotype. Its results demonstrate a male-specific time-related loss of nigral dopamine neurons, reductions in striatal dopamine levels, and motor dysfunction and elevation of alpha-synuclein. As such, these findings are critical not only to understanding the contribution of PQ exposure to the etiology of PD, but ultimately to the regulation of PQ usage in the US to safeguard public health.

## RESULTS

### PQ Exposure Characteristics

**Figure 1** presents PQ exposure characteristics across exposure sessions. As shown, the average particle count over the 60-day exposure was 24,438 ± 5,355 particles/cm^3^ (**Figure 1A**), compared with 21,900 ± 3,320 particles/cm^3^ from the previously reported 20-day exposure paradigm [17]. The average particle mass concentration (**Figure 1B**) across exposure was 122 ± 38 μg/m^3^, and median particle diameter (**Figure 1C**) was 198 ± 6.67 nm (1.46 Geo. SD), smaller than the typical size range reported in agriculture by the EPA (400-800 μm) [21]. Our previous inhalation study produced an average particle mass concentration of 130 ± 10 µg/m^3^, and an average particle diameter of 166 ± 3.1 nm (1.43 Geo. SD) across 20 days of PQ exposure [17]. Thus, exposure metrics were all within ∼10-20% of those from the previous study, demonstrating the reproducibility of this inhalation model.

**Figure 1.**
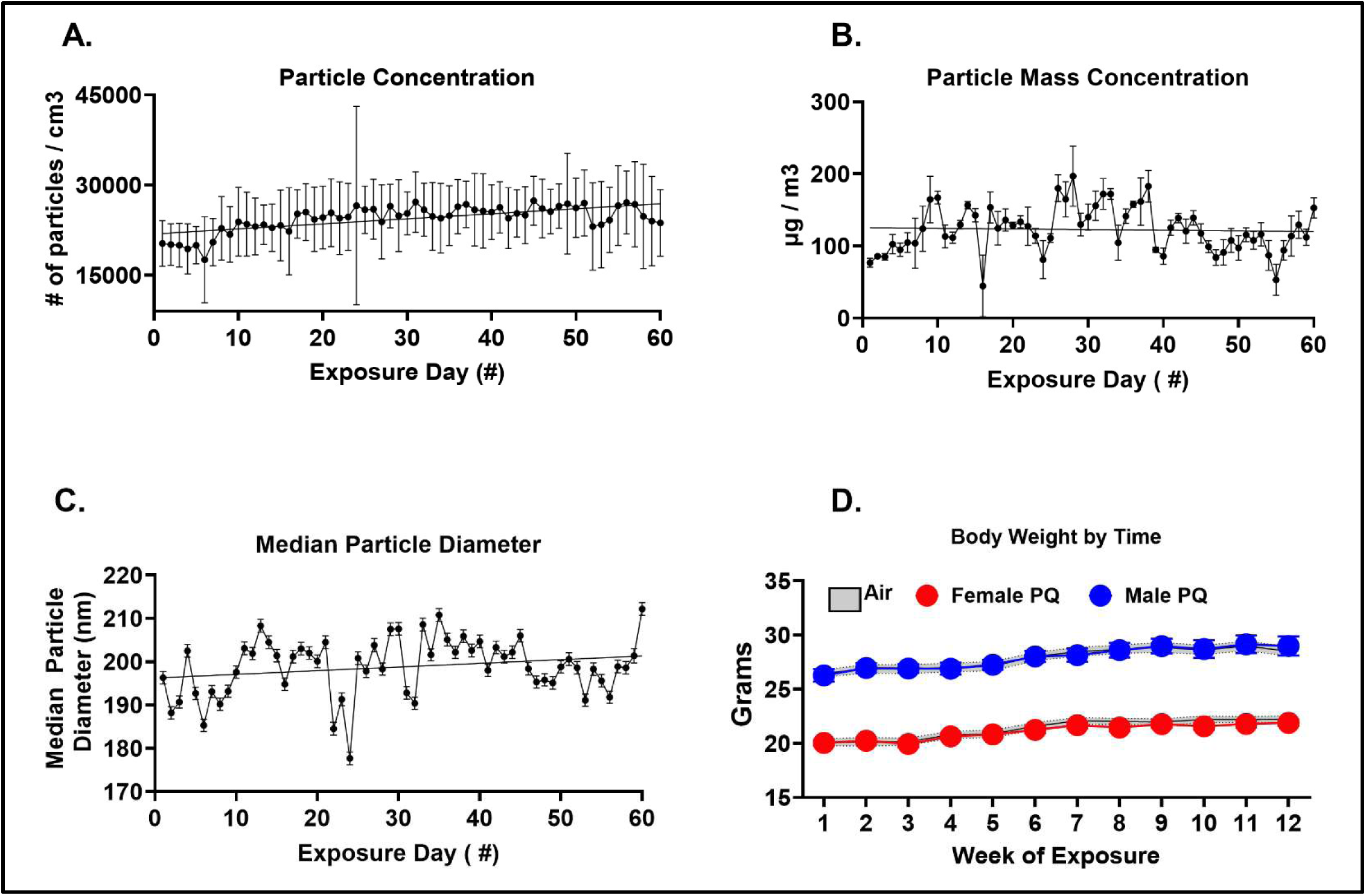
PQ Exposure Characteristics and Body Weights. Exposure conditions for 60-day PQ inhalation treatment (5-days/week, 4 hours/day), including (A) measurements of PQ particle concentration (B) particle mass concentration and (C) mean particle diameter. (D) Body weights (grams) in female and male mice exposed to filtered air or PQ for 3 mos; statistical significance determined by Two-Way ANOVA with Tukey’s multiple comparison test.

### PQ and Body Weight Across Exposures

PQ exposures did not result in any changes in body weight (**Figure 1D**) as assessed weekly in the 3 mos cohort, in either females or males.

### PQ Uptake into Brain Following Inhalation Exposures

In the brain, the highest levels of PQ (**Figure 2**) were found in the olfactory bulb and cortex in both sexes. In the case of the olfactory bulb, increases were evident after 2 mos of PQ exposure in females (F(1,3)=49.68, p=0.0059), with a further significant increase of approximately 215% relative to the 2 mos time point seen after 3 mos of PQ exposure (F(1,3)=1183.8, p<0.0001). Males showed a similar pattern with nearly significant increases after 2 mos of PQ exposure (F(1,4)=7.51, p=0.052) with an approximately further increase of 102% in PQ levels between 2 and 3 mos of exposure (F(1,3)=17.43, p=0.025). In the case of the frontal cortex, significant increases in PQ levels were evident after 2 mos of exposure (F(1,4)=8.75, p=0.0416). with a further increase of approximately 263% after 3 mos of exposure (F(1,4)=24.13, p=0.008). The 2 mos PQ exposure produced an increase in levels in the frontal cortex (>2000%) that was marginally significant (F(1,4)=6.87, p=0.0588) with a further increase of 287% after 3 mos of PQ exposure (F(1,4)=4.42, p=0.1035), albeit not statistically significant. Of the three regions examined, the lowest levels of PQ in the brain were found in the brainstem. Nevertheless, levels of PQ were increased by more than a thousandfold after 2 mos of exposure, a marginally significant increase (F(1,4)=7.56, p=0.051) in females, with a further increase of 140% after 3 mos of exposure (F(1,4)=428.34, p<0.0001. In males, a marginal increase was found after 2 mos of PQ exposure (F(1,4)=5.54, p=0.078, which slightly declined after 3 mos of exposure but was nevertheless still significantly increased (F(1,3)=18.69, p=0.0228).

**Figure 2.**
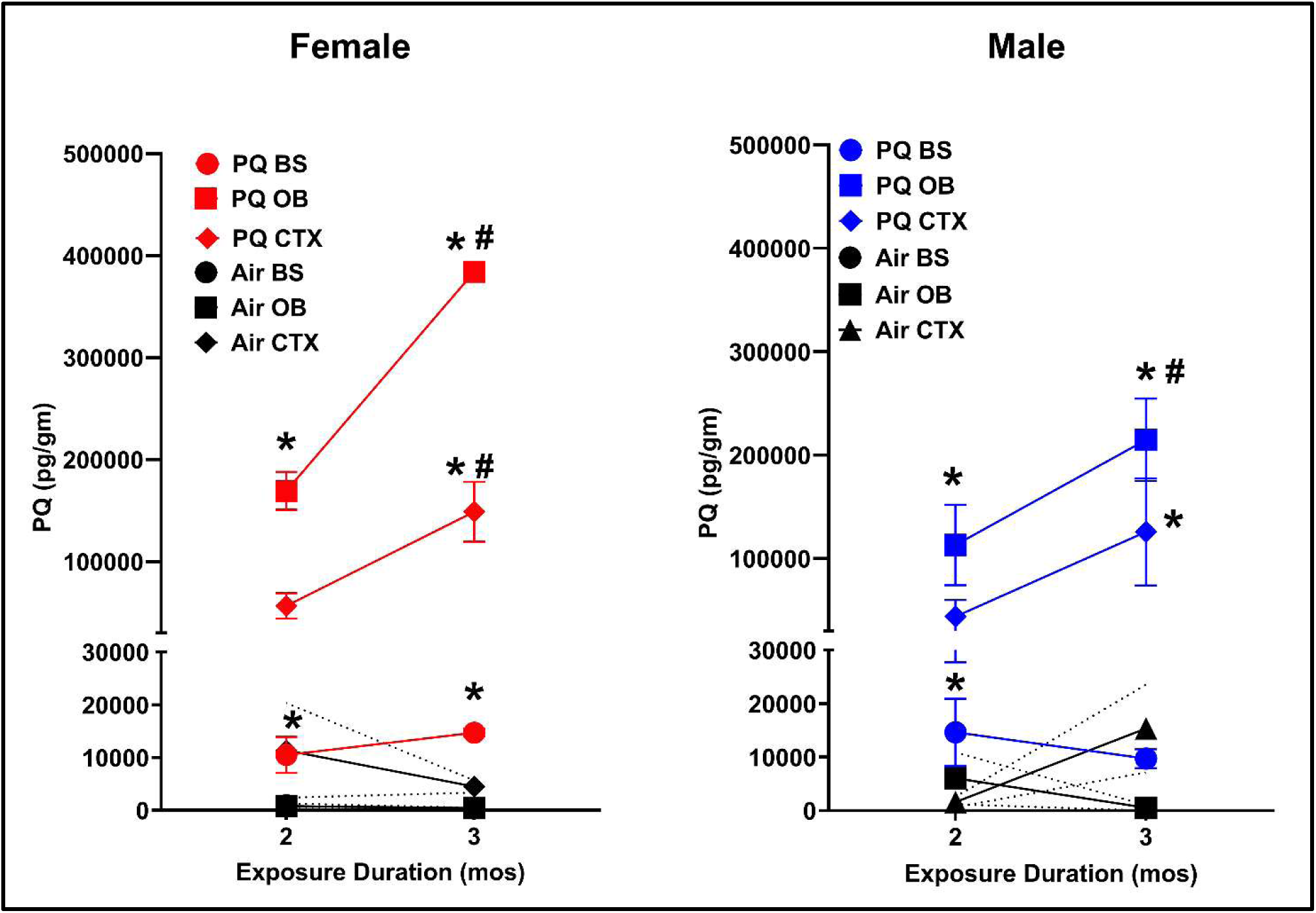
PQ Quantification in Olfactory Bulb, Cortex, and Brainstem. Group mean ± standard error levels of PQ (picogram/gram) in the hemisected olfactory bulb, frontal cortex, and brainstem (as indicated) in female (left panel) and male (right panel) mice exposed to inhaled PQ for 2 or 3 mos, as indicated. n = 3/sex/treatment group/time point. Statistically significant differences were determined by Two-Way ANOVA with Tukey’s multiple comparisons test. *= significantly different from 3 mos filtered-air control and 2 mos PQ and 2 mos filtered-air control values; #= significantly different from 2 mos filtered-air control value. Sample sizes were n =3 per sex/treatment group per time point.

### PQ Inhalation Induced Male-Specific Reductions in Locomotor Activity

To determine whether PQ inhalation leads to a Parkinsonian motor phenotype in mice, a behavioral battery of motor endpoints was carried out following both the 2 mos (40 exposure days) and 3 mos (60 exposure days) exposure time points. It revealed male-specific reductions in locomotor activity (**Fig. 3 top row**) ranging from 23-27% after the 3 mos PQ exposure relative to filtered air controls, which were not yet seen after 2 mos of PQ exposure. These changes consisted of significant reductions in ambulatory metrics in the center compartment, including time, counts, and distance (time x treatment p=0.0136, 0.0189, and 0.0151, respectively). Additionally, marginal reductions were observed in the center compartment for both stereotypic time and stereotypic counts (time x treatment p=0.0763 and 0.066, respectively). Females, in contrast to males, showed marginal increases in ambulatory activity metrics at the 2 mos time point (F(1,14)= 3.62, p=0.078, 4.08, p=0.063, and 3.46, p=0.084 for ambulatory time, counts and distance, respectively).

**Figure 3.**
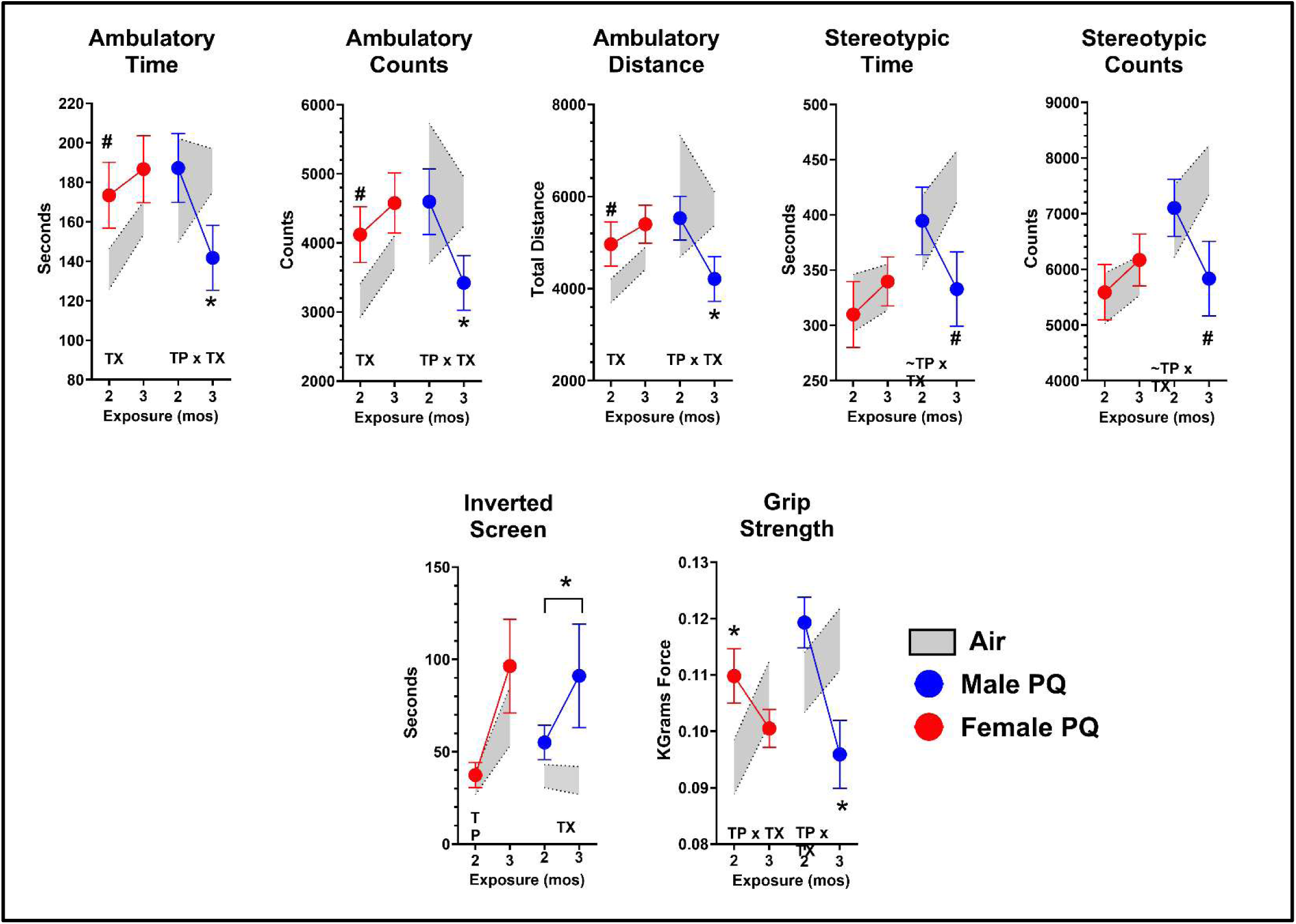
Male-Specific Motor Deficits in PQ-Exposed Mice. Group mean ± standard error changes in measures of locomotor activity, including, from left to right, ambulatory time, ambulatory counts, ambulatory distance, stereotypic time, and stereotypic counts in PQ-exposed male and female (as indicated) mice relative to filtered-air control mice. TX = p<0.05; main effect of treatment (PQ). TP x TX = p<0.05; interaction of time point by treatment. Statistical significance was determined by a Two-Way ANOVA with Tukey’s multiple comparisons test. *= p<0.05; significantly different from corresponding filtered-air control values. #= p<0.06; marginally significant from corresponding filtered-air controls. Sample sizes of n = 8 per sex/treatment group/time point.

Males exposed to PQ for 3 mos likewise showed deficits in both inverted screen performance and grip strength (**Figure 3, bottom row**). Significant increases in time-to-right on the inverted screen appeared in response to both the 2 and 3 mos PQ exposures (F(3,27)=3.1, p=0.043, main effect of treatment, p=0.0154). The PQ-exposed males took 50% longer to right themselves than filtered-air controls at the 2 mos time point and 265% longer after 3 mos of PQ exposure. While females actually exhibited a marginally significant increase in grip strength at the 2 mos time point (F(3,28)=2.35, p=0.096, time x treatment, p=0.0234), reductions of approximately 20% were evident in PQ-exposed males at the 3 mos time point (F(3,27)=3.62, p=0.026, time x treatment, p=0.0075) relative to male filtered-air controls.

### PQ Inhalation Altered Brain Cysteine Pathways

Given the potential redox capacity of PQ [22], an assessment of the brain transsulfuration pathway was performed in the striatum (**Figure 4)**. These measures revealed interactions between PQ and the cysteine pathways in both sexes. Females showed marginal increases in striatal cysteine after 2 mos of PQ exposure (F, 3,20)=3.11, p=0.0493; treatment by time, p= 0.0590) with a trend towards subsequent reduction at 3 mos. While males showed a similar overall pattern, statistically, analyses revealed a significant time-by-treatment effect (F(3,20)=4.95, p=0.0099; time x treatment p=0.0155), including a marked 67% reduction after 3 mos of PQ exposure. No significant changes in methionine levels or in glutathione were seen in either sex.

**Figure 4.**
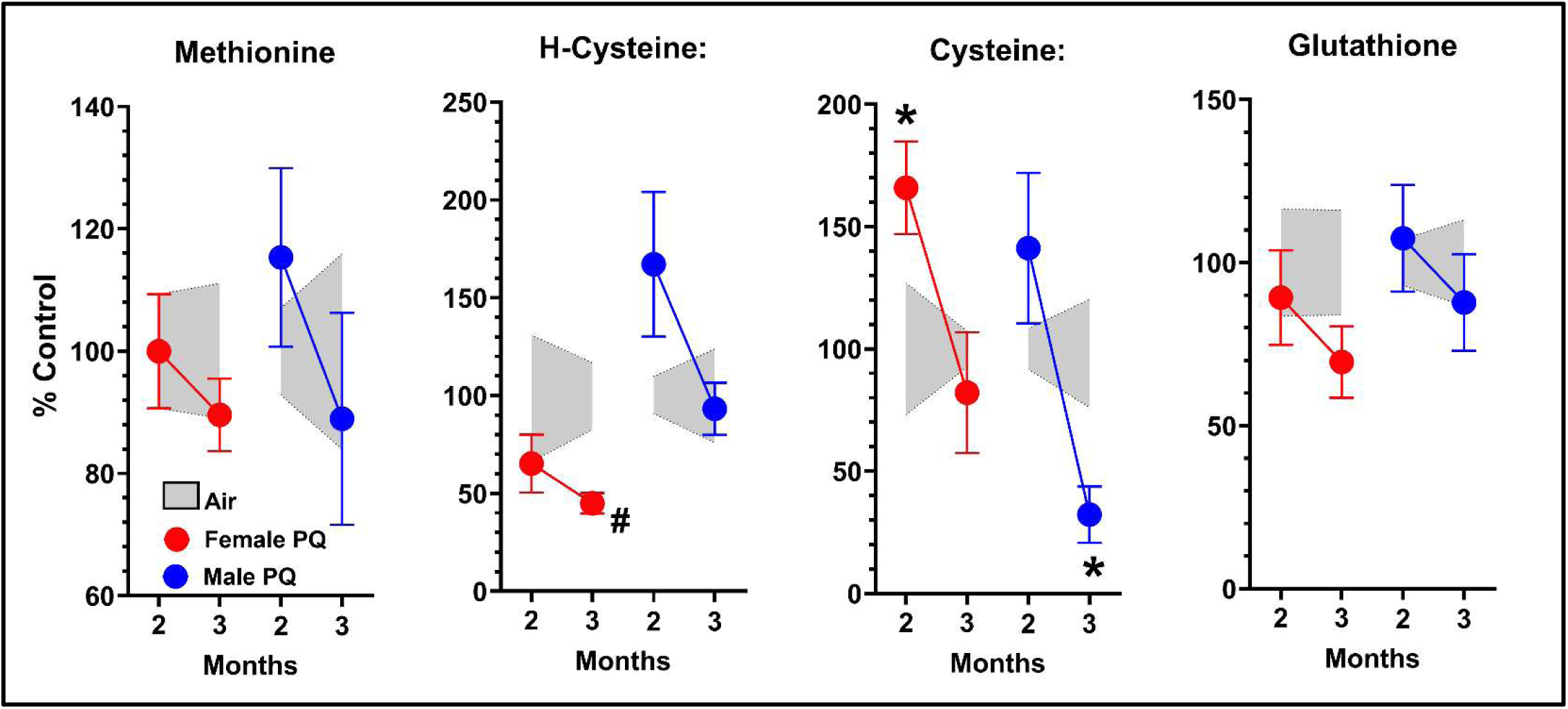
PQ-induced Changes in Transsulfuration Pathway Across Sexes. Group mean ± standard error levels of striatal methionine, homo-cysteine, cysteine, and glutathione in female and male (as indicated) PQ-exposed mice relative to filtered-air control values (as indicated) at the 2 and 3 mos time points. Statistically significant differences were determined by using Welch’s t-test. *= p<0.05; significantly different from corresponding filtered-air control values. Sample sizes of n = 6 per sex/treatment group/time point.

### PQ Inhalation Produced Male-Specific Increases in Total α-Synuclein Levels in the Olfactory Bulb

Accumulation of misfolded α-synuclein represents a hallmark of Parkinson’s disease [23] and both extra-nuclear and nuclear levels were measured in both the olfactory bulb and the substantia nigra pars compacta (SNc), as shown in **Figure 5**. Analyses revealed male-specific increases in total extra-nuclear α-synuclein levels in response to 3 mos of PQ exposure that were seen in the olfactory bulb, but not in the SNc, and were not seen in nuclei in either region. Specifically, levels of total α-synuclein in 3 mos PQ-exposed males were increased approximately 85% above filtered air control values (F(3,19)=5.90, p=0.005; time x treatment, p=0.0244) while the total synuclein area of 3 mos PQ-exposed males increased by 70% (F(3,19)=5.44, p=0.0072, time x treatment, p=0.0244).

**Figure 5.**
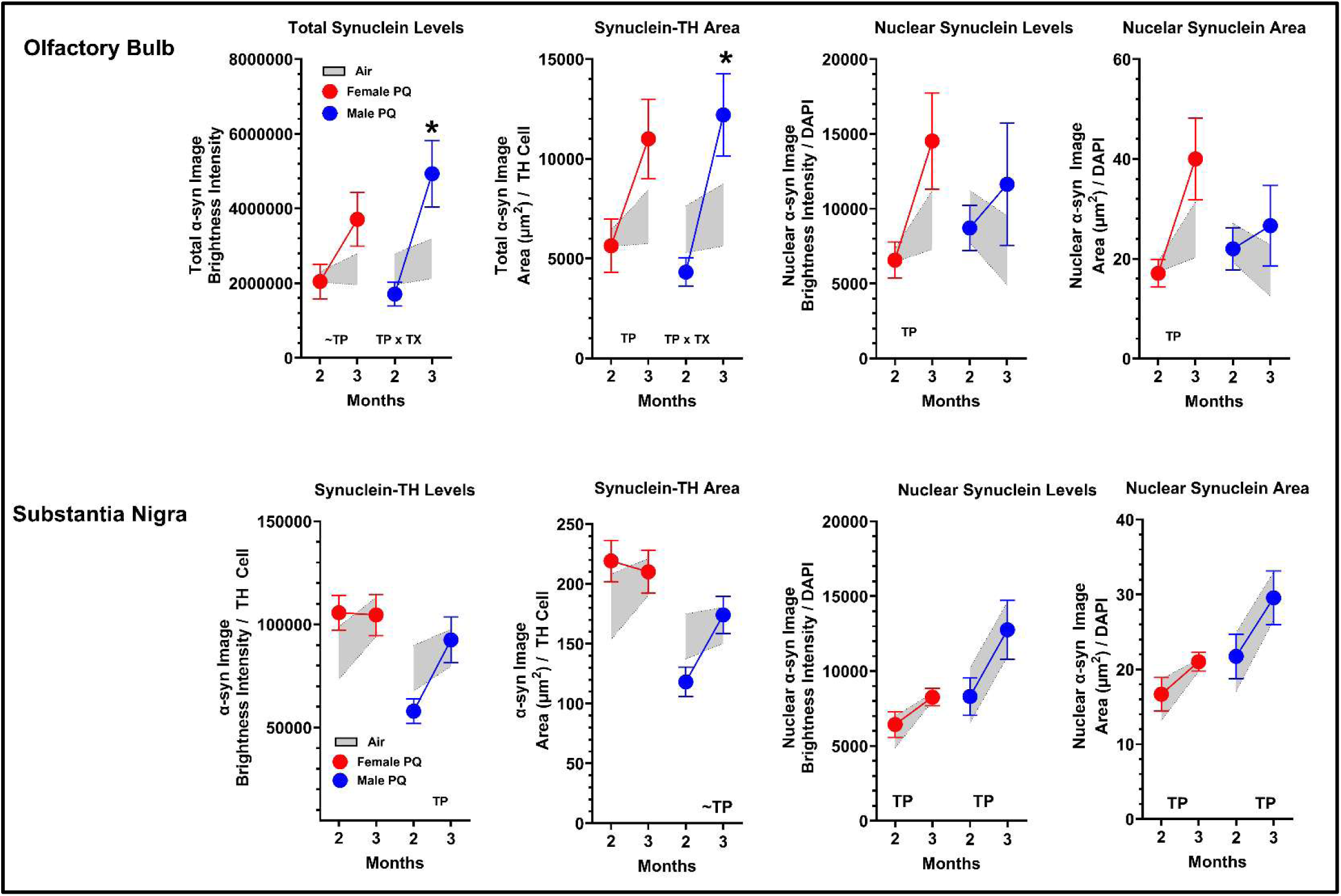
Male-Biased Increase in Total α-Synuclein in the Olfactory Bulb and Substantia Nigra. Group mean ± standard error changes in olfactory bulb (top row) and substantia nigra (bottom row) of total α-synuclein levels and area, and nuclear α-synuclein levels and area from left to right in each row in female and male (as indicated) PQ-exposed mice relative to filtered air control. Statistically significant differences were determined by Two-Way ANOVA with Tukey’s multiple comparisons test. TP = p<0.05; main effect of time point. TX = p<0.05; main effect of treatment (paraquat). TP x TX = p<0.05; interaction of treatment by time point. *= p<0.05; significantly different from filtered air control. Sample sizes were n = 6 per sex/treatment group/time point.

### PQ Inhalation Produced Male-Specific Reductions in Levels of Striatal Dopamine

One of the key hallmarks of PD is the loss of dopaminergic neurons, which express the rate-limiting enzyme for DA synthesis in the SNc, tyrosine hydroxylase (TH) [24, 25]. The progressive loss of these neurons leads to reduced dopamine (DA) biosynthesis, a neurotransmitter critical for various functions, including voluntary movement. Many of the locomotor symptoms of PD begin to manifest once a critical mass of dopaminergic neurons in the SNc die or degrade, where the remaining population can no longer compensate for the body’s demand for DA production, and the labile pool is significantly reduced [26, 27]. To investigate immediate biochemical changes induced by PQ inhalation, changes in brain neurotransmitter levels were assessed in the striatum of both sexes immediately following each time point.

These analyses revealed that while markers of glutamatergic and serotonergic neurotransmitters were largely unaffected, marked striatal dopaminergic dysfunction was seen in PQ-exposed males at the 3 mos time point (**Figure 6**). This included reductions in 3,4-dihydroxyphenylacetic acid (DOPAC), the first metabolite in the DA degradation pathway produced by monoamine oxidase (MAO), of 63% (F(3,20)=2.63, p=0.079; main effect of treatment, p=0.0.0188). In addition, a 73% reduction in DA was observed (F3,20)=4.44, p=0.0151, main effect of treatment, p=0.0118). This was accompanied by marked increases of >1000% in measures of DA synthesis as indicated by the ratio of the precursor tyrosine to DA (F(3,20)=3.86, p=0.0251; main effect of treatment, p=0.0355) with marginal increases in markers of DA turnover, including both the ratio of DOPAC/DA (F(3,20)=2.49, p=0.09) and of homovanillic acid (HVA)/DA (F(3,20)=2.38, p=0.100).

**Figure 6.**
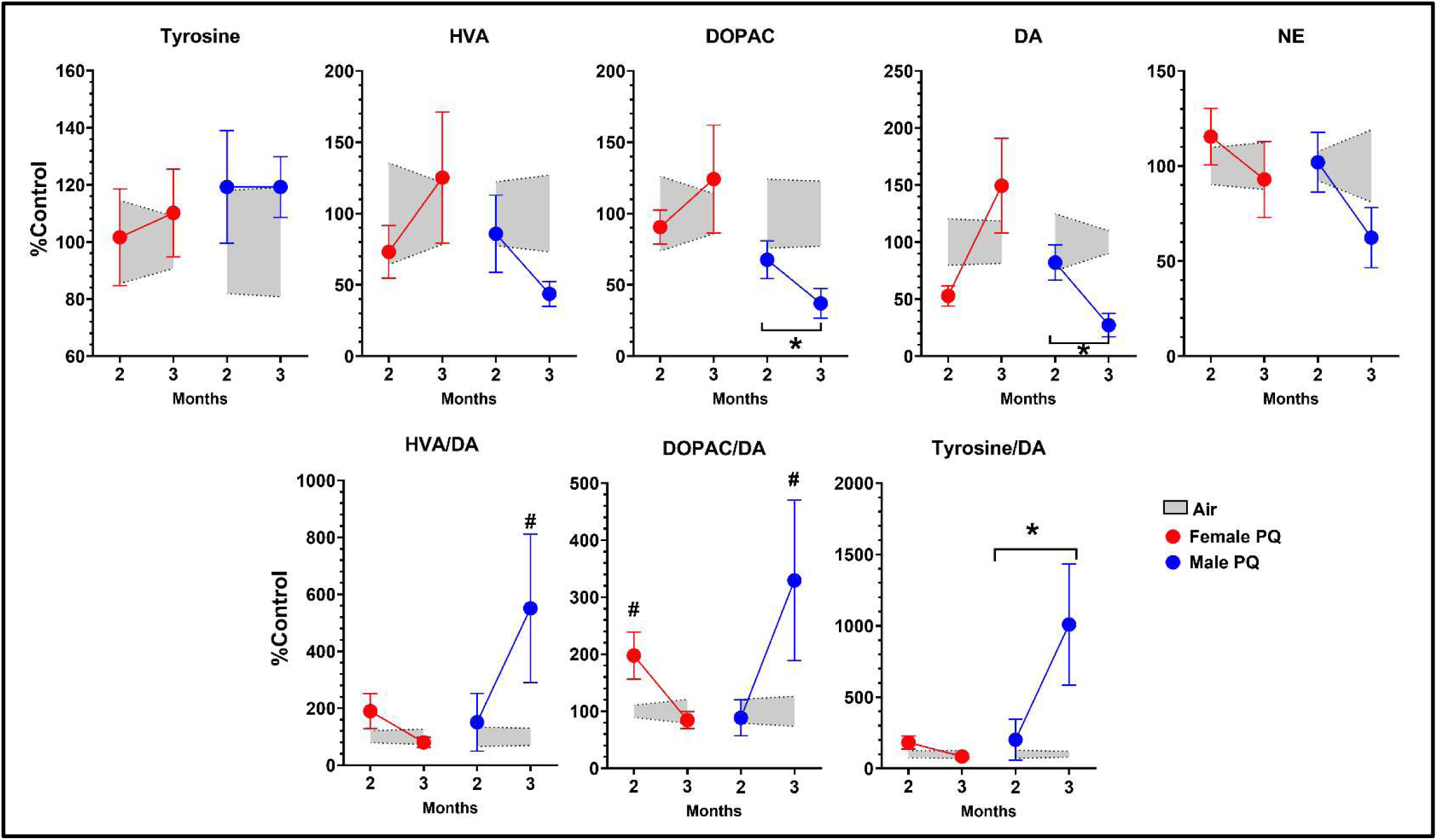
PQ Inhalation Induces Male-Biased Reduction in Striatal Dopaminergic Neurotransmitters and Turnover After 3 Months. Group mean ± standard error of dopaminergic neurotransmitters and turnover in female and male (as indicated) mice exposed to PQ for 2 or 3 mos plotted as percent of filtered-air control values. Statistically significant differences were determined by Two-Way ANOVA with Tukey’s multiple comparisons test. *= p<0.05; significant effect of PQ treatment relative to filtered-air control at both time points. #= p<0.05; significant effect of PQ relative to filtered-air control at 3 mos time point. Sample sizes were n =6 per sex/treatment/time point.

The only effect observed in females was an increase in the DOPAC/DA ratio (F3,20)=4.33, p=0.0165; effect of time by treatment, p=0.035) following 2 mos of PQ exposure. However, this effect was no longer observed at the 3 mos time point.

### PQ Inhalation Produced Male-Specific Reductions in Tyrosine Hydroxylase (TH) Levels in Substantia Nigra

Levels of TH, the enzyme that catalyzes the formation of L-DOPA (the rate-limiting step of DA biosynthesis), were measured here in both the olfactory bulb and striatum (**Figure 7**). As can be seen, male-specific increases in TH levels were seen only in PQ-exposed males and in the olfactory bulb but not in the SNc that were statistically marginal (F(3,19)=2.01, p=0.146, main effect of treatment p=0.0248 and F(3,19)=1.81, p=0.18, main effect of treatment, p=0.0311). Increases in TH level averaged 3% after 2 mos PQ exposure and 11% after 3 months of PQ exposure, with corresponding values of 4% and 10% for TH area.

**Figure 7.**
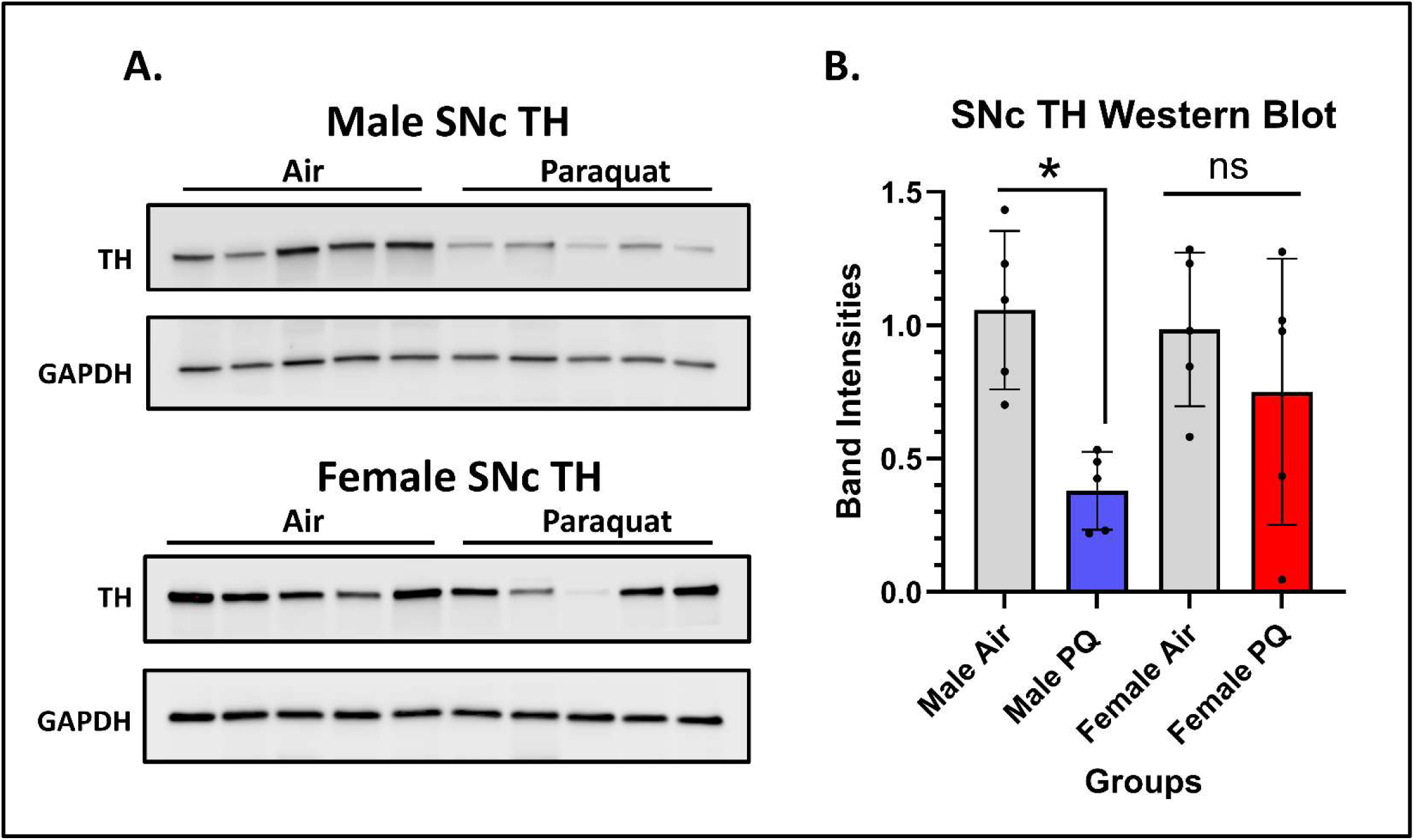
Male-biased Reductions in Tyrosine Hydroxylase Expression in Substantia Nigra After 3 Months of PQ Inhalation. **Panel A**. Blots of TH and GAPDH in Air-exposed and paraquat-exposed male mice. **Panel B.** Group mean ± standard error band intensities in the 3 mos cohort of filtered-air controls and PQ exposed males and females as indicated. Statistical significance was determined using Welch’s t-test. *= p<0.05; significant effect of PQ treatment relative to filtered-air controls. Sample sizes were n = 5 per sex/treatment.

### PQ Inhalation Produced Male-Specific Reductions in Numbers of Tyrosine Hydroxylase Positive Neurons in Substantia Nigra

The loss of dopaminergic neurons in the SNc is known to underlie the motor symptomatology of PD. In the current study, the number of TH+ neurons in the SNc was assessed after both the 2 and 3 mos exposure timepoints. Results confirmed male-biased reductions in the number of SNc TH+ dopaminergic neurons observed after 3 mos of PQ inhalation exposure **(Figure 8)** averaging approximately 17% ((F(1,8)=6.84, p=0.031), but were not yet present at 2 mos of PQ exposure. No changes were found in females.

**Figure 8.**
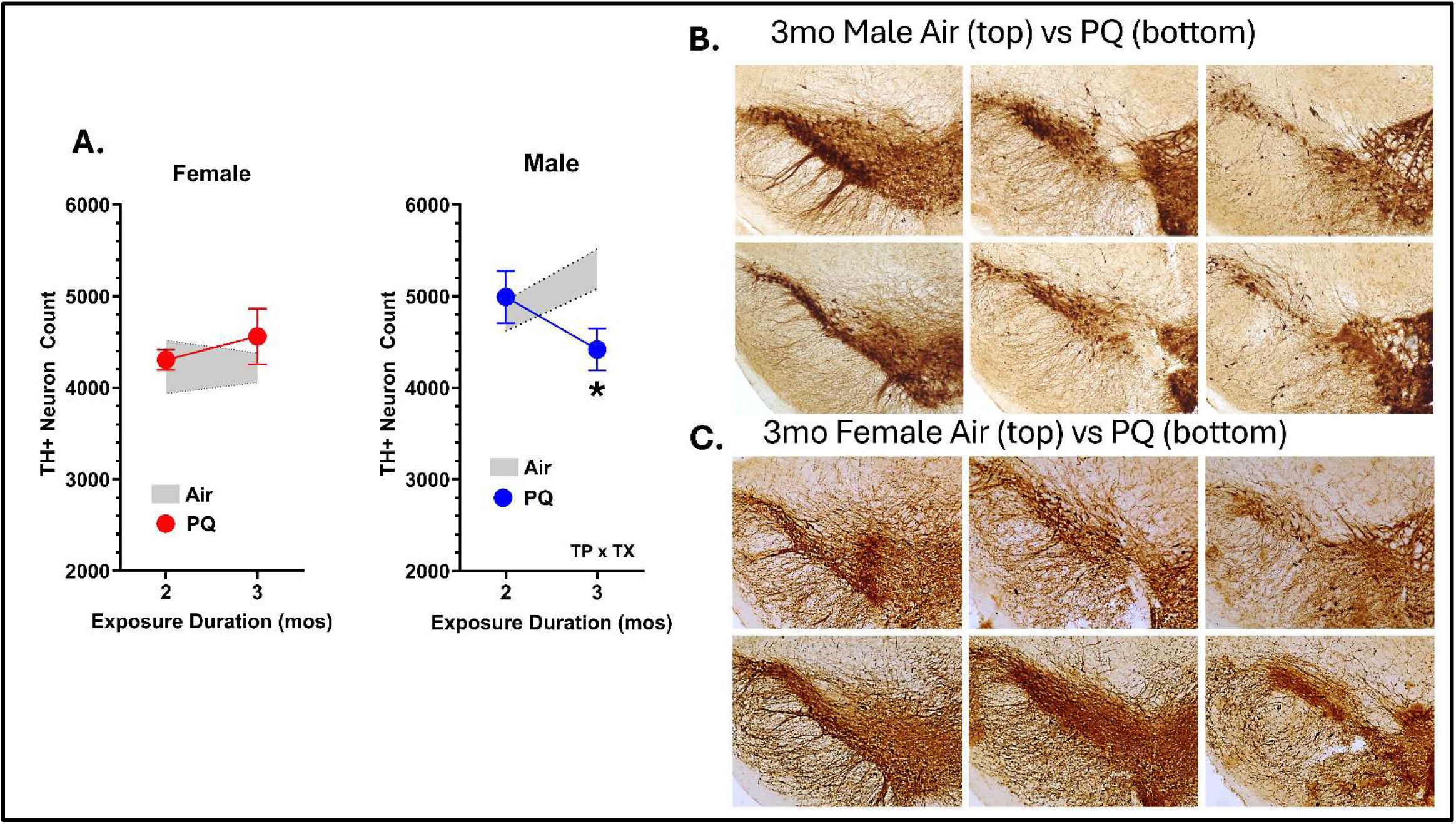
Male-Biased Reductions in TH+ Dopaminergic Neuron Density in the Substantia Nigra Following 3 Months of PQ Inhalation. **A.** Group mean ± standard error changes in TH+ neuron density in substantia nigra in females (left) and male (right) mice exposed to PQ or filtered-air at the 2 and 3 mos time point. Statistically significant differences were determined by Two-Way ANOVA with Tukey’s multiple comparisons test. TP x TX = p<0.05; interaction of treatment by time point; *= p<0.05; significantly different from filtered-air control. Sample sizes were n = 6 per sex/treatment group. **B.** Representative images of TH+ neuron density in male substantia nigra after 3 mos exposure to filtered-air (top row) or PQ (bottom row). **C.** Representative images of TH+ neuron density in female substantia nigra after 3 mos exposure to filtered air (top row) or PQ (bottom row).

## DISCUSSION

Parkinson’s disease (PD) is the fastest-growing neurodegenerative disease in the world [1, 28], of which only a small fraction of cases, estimated to be 10% or less, are attributable to the inheritance of a monogenic mutation alone [29, 30]. In fact, most genetic variants are low-penetrance risk factors that do not directly cause PD [31, 32]. Such findings underscore the importance of environmental risk factors for PD. One such risk factor that has repeatedly been associated with an increased risk of PD, both in epidemiologic studies [33–35] and animal models [36, 37] is exposure to the herbicide paraquat (PQ). However, the model system approaches have been criticized for using systemic PQ exposures, e.g., intraperitoneal (IP) injections, that do not mimic human exposures, which are largely considered to result primarily from inhalation exposures but can also include a dermal component [38].

In a prior study examining a 20-day paradigm of PQ inhalation, a male-specific olfactory impairment was observed, consistent with prodromal features of PD. The aim of the current study was to assess the impact in mice of more protracted PQ inhalation exposures, i.e., 40 days and 60 days, with respect to reproducing characteristics of PD. This included an assessment of motor function, levels of striatal neurotransmitters and dopaminergic function, levels of tyrosine hydroxylase (TH), and of α-synuclein and the estimated density of TH+ neurons in the substantia nigra pars compacta (SNc), all known components of PD [39–42]. Collectively, the findings show that inhalation of PQ produced male-specific features of PD that were duration-of-exposure-dependent, emerging after 60 days (3 mos) of exposure. These included reductions in motor function, reductions in striatal dopamine (DA), increases in total α-synuclein levels in the olfactory bulb, alterations in homocysteine/cysteine pathways, and reductions in TH expression in the SNc and in the number of SNc TH+ neurons, all findings consistent with PD pathophysiology, including the male-bias of effects. Collectively, these findings clearly support the potential for prolonged PQ inhalation to induce hallmarks of the PD phenotype.

Levels of PQ were found to increase in the olfactory bulb and frontal cortex between the 2 and 3 mos exposure period in both sexes, findings consistent with cumulative exposures, whereas lower, more consistent levels of PQ were found in the brainstem. These results demonstrate that inhaled PQ does reach the brain. They are also not inconsistent with findings reported regarding the toxicokinetics of PQ in the brain following systemic exposures. Specifically, Prasad et al. [43] reported a half-life of approximately 28 days following IP injections of 10 mg/kg PQ in mice. Similarly, another study of PQ using IP injection reported a slow elimination of PQ from the brain, citing a 21-day half-life [44]. In our prior study using a 20-day total PQ inhalation exposure, PQ levels in the olfactory bulb were negligible when measured 28 days post-exposure [16]; however, when levels were determined as PQ per region/organ, the olfactory bulb and brainstem had higher burdens than other regions [16]. Additional studies to define the toxicokinetics in the brain following inhalation exposure could help further elucidate the time course of uptake and elimination of inhaled PQ.

Male-specific impairments in motor function were observed after 3 mos of PQ inhalation exposure, as assessed in an open field, with reductions in ambulatory-related activity and distance, as well as decreases in grip strength and increased time to righting on an inverted screen. Correspondingly, systemic exposures to PQ in rodent models have reported alterations in motor function, including reductions in locomotor activity following IP exposures [45] as well as in response to subcutaneous exposures [46].

In addition to the emergence of motor deficits following 3 months of PQ inhalation, increases approximating 70-85% in total α-synuclein levels, a hallmark of PD, were observed in the olfactory bulb in males, but were not evident in the SNc. Increases in total α-synuclein in the brain have similarly been reported in response to systemic PQ exposures, including in regions such as the ventral mesencephalon and frontal cortex [47]. That dose levels and timing are important to the ultimate impact of PQ exposure on α-synuclein accumulation was indicated by reports that a systemic 10 mg/kg IP injection administered once a week for 4 weeks, rather than the typical 3 weeks, led to reductions in total α-synuclein expression in the SNc, but marked increases in hippocampal levels [48]. Increases in pSer129 α-synuclein have also been reported in response to 50 ug/ml PQ *ad libitum* oral exposures for 6 weeks in the TgM83 (HuA53T-α-syn) transgenic mouse model [49].

Increases in α-synuclein levels are not inherently pathological, given that the protein is natively expressed in the CNS, where it is believed to aid presynaptic vesicle trafficking and exocytosis [50]. These increases in total α-synuclein levels may predict future misfolding, as upregulation is a concentration-dependent driver of aggregation [51]. Phosphorylation at Ser129 of α-synuclein is a more specific marker of pathological aggregates, but recent studies have shown that they are also present in healthy mammalian brains [52]. Differentiating between healthy and pathogenic pools of pSer129 α-synuclein is necessary for precise quantification of pathological inclusions. Therefore, further studies should focus on aggregated α-synuclein levels to assist in refining this biomarker of PD.

Male-specific reductions in striatal DA levels, SNc TH expression levels, and numbers of TH+ neurons emerged after 3 mos of PQ inhalation, with the TH+ neuron loss averaging approximately 17%, indicating an as yet mild Parkinsonian phenotype. To the authors’ knowledge, this is the first study to show dopaminergic neurodegeneration in the SNc following PQ inhalation. Loss of TH+ neurons following inhalation exposures has also been reported after systemic PQ administration [53–56]. Mice in the current study were not aged following their initial exposure and subsequent behavioral assessment. It is possible that assessments over a longer post-exposure period would reveal more acute locomotor dysfunction or a greater magnitude of TH+ neuron loss. Consequently, future studies using this model and testing the hypothesis that exposure effects worsen over time with age after inhalation exposure ends would be informative.

Another interesting tie between PQ inhalation was the disruption of the homocysteine/cysteine and transsulfuration network in the striatum, another marker of adverse effects in PD and an important regulator of dopaminergic cell death and glutamatergic function [57]. Specifically, these were seen as increases in cysteine after 2 mos of PQ exposure and reductions in homocysteine after 3 mos of PQ exposure in females, while males showed trends toward increased homocysteine and cysteine at 2 mos, with a sharp reduction in levels of cysteine after 3 mos of PQ exposure. Changes in homocysteine have been implicated in increased inflammation and oxidative stress, with consequences that can include brain atrophy, apoptosis, and excitotoxicity [58]. Increases in plasma/serum total homocysteine have been reported in PD and associated with dementia [59], exacerbation of motor deficits and dopaminergic neurodegeneration [60]. The reduction in brain cysteine levels in males is notable, given its role as the rate-limiting step in the synthesis of the major antioxidant glutathione [61]. While no reductions were seen at this time in striatal glutathione levels, and the reduction in dopaminergic neurons in the SNc is mild following the 3 mos PQ exposures in males, a more detailed time course of the transsulfuration pathway changes will be informative in understanding the trajectory of effects and their ultimate consequences.

Very little information exists regarding exposures to airborne pesticides in the US, likely due to the inherent variability of environmental factors such as temperature, humidity, and wind speed, which make consistent measurements difficult [62]. A previous field study measuring PQ concentrations near treated cotton fields measured 4.31 to 10.7 μg/m^3^ at 1 meter downwind and less than 50 ng/m^3^ at 400 meters [63]. A similar study measuring occupational exposure to inhaled PQ measured an average daily exposure for field workers of 4.89 ± 10.77 μg/m^3^, with the highest single-day exposure being 93.12 μg/m^3^ [64]. The exposure concentration used in our study, 122 ± 38 μg/m^3^, is closest to this peak single-day exposure, accounting for variability across these field studies and reports indicating that pesticide loaders, mixers, and workers are exposed to higher concentrations of airborne pesticides [65].

Male-bias is a well-known component of PD [66]. As noted, the effects of PQ inhalation in this study were primarily male-specific, with PQ-exposed females showing no consistent changes in motor function, striatal DA, or changes in TH+ neuron counts and TH expression in the SNc. Male susceptibility to PQ is not limited to murine models but has also been seen in rats [67] and *Drosophila* models [68, 69] of PQ neurotoxicity. These results are consistent with the literature, as neurotoxicology studies have historically found male-specific vulnerabilities to neurotoxicants associated with PD [70]. For example, studies of rotenone (IP) [71], dieldrin (dietary) [72], and manganese (subcutaneous) [73] exposure in rodents have observed male-specific susceptibility to toxicant-induced α-synuclein aggregation, medium spiny neuron spine density, and TH+ neuron loss in the SNc. The reason(s) for the male vulnerability and the resistance of females remain unknown. Numerous lines of evidence also suggest the potential role of estrogen (estradiol in mice) as a putative protective factor for dopaminergic dysfunction in females. Estrogen has been shown to increase the synthesis, release, reuptake, and turnover of DA [74]. In addition, estrogen shows anti-inflammatory properties [66]. In addition to hormonal differences, sex differences in neuroimmune function [75] and numerous differences in SNc function and structure, including differences in lipids [76], and in dopaminergic and cholinergic activity and function [77].

In addition to hormonal differences, emerging evidence has shown that sex-determining region of the Y chromosome (SRY) may play a role in male susceptibility to neurotoxicants. SRY is present in a subset of midbrain dopaminergic neurons in males [78], and serves as a transcription factor for dopamine-related enzymes (TH, monoamine oxidase-A, and DA decarboxylase) [79]. In rats, inhibition of SRY in males protected against rotenone and 6-hydroxydopamine-induced nigrostriatal neurodegeneration [80], suggesting its expression contributes to male susceptibility to neurotoxicants. SRY overexpression has also been shown to reduce expression of the estrogen receptor beta (ERβ) in mouse Neuro 2A cells, which may hamper estrogen’s neuroprotective effects in males [79]. Therefore, male susceptibility to PD-associated neurotoxicants is multifaceted and influenced by factors upstream of hormonal signaling.

Collectively, the data in this study are consistent with an olfactory bulb portal of entry for PQ into the brain with subsequent transport to the frontal cortex. This may occur by circumventing the BBB and entering the CNS through the primary sensory neurons in the olfactory epithelium, which are directly exposed to the environment [81, 82]. Despite being charged, hydrophilic ions, PQ can also cross the BBB through a commonly expressed family of neutral amino acid transporters [83], offering an alternative mode of transport if PQ aerosols are partially ingested. The far smaller increases in PQ levels in the brain stem as compared to the olfactory bulb and frontal cortex suggest either slower redistribution from the olfactory cortex and/or no real vagal transport into the brain. The second-highest concentration of PQ in the brain was found in the cortex, followed by the brainstem, indicating that PQ can readily distribute throughout the brain prior to clearance.

The current study extends the mouse model of low-dose PQ inhalation first reported in Anderson et al. [16], from an initial 20-day exposure to 40 or 60 days, to assess the progression of biochemical and neurobehavioral changes associated with dopaminergic circuitry. Collectively, findings from the current study and Anderson et al. [16] unveil a trajectory of changes consistent with prodromal PD, e.g., male-specific impaired olfactory discrimination [16] seen after a 20-day PQ inhalation exposure, followed by emergence of increased total α-synuclein levels in the olfactory bulb, reductions in striatal DA, in levels of TH expression and TH+ neuron count in the SNc, and impaired motor function after 3 mos of exposure. Notably, the effects of PQ inhalation exposure shown here replicate those reported following systemic exposure in rodent models [47, 53–56, 84]. This suggests that, regardless of the route of exposure, once PQ enters the brain, it produces Parkinsonian features, indicating its potential health hazard. Further studies of PQ inhalation exposure, including exposures earlier in the life span, will be important for understanding the cumulative impact, the potential to prime the brain for subsequent vulnerability, and ascertaining the mechanisms underlying its effects.

## METHODS

### Animals

Male and female C57BL/6J mice (Jackson Laboratories) pair-housed in standard mouse caging with ∼3 mm high-performance bedding (BioFresh) and supplied with standard rodent chow (LabDiet Autoclavable Diet 5010) and water *ad libitum* were maintained at 22°C ± 2°C under a 12-hour light-dark cycle at the University of Rochester Medical Center. At postnatal week 14 (adulthood), mice were randomly assigned to either a PQ or filtered-air control group. Within each treatment group, mice were further divided into either a behavior or non-behavior group, as behavioral assessments have the potential to modify the effects of neurotoxicants [85]. Sample sizes were n = 7-8/treatment/behavior group for males and females.

To determine potential PQ deposition in brain and assess markers of neurodegeneration, a subset of mice was sacrificed immediately after exposure ended at 2 and 3 months. The behavior group underwent behavioral testing and were sacrificed two weeks post-exposure. Mice were weighed every Friday, throughout the study, to assess signs of systemic toxicity and stress. All mice were treated humanely and with regard for alleviating suffering. Mice that presented with consistent weight loss and/or signs of excessive pain or discomfort were euthanized to minimize suffering according to University policy. All procedures were approved by the Institutional Animal Care and Use Committees at the University of Rochester, i.e., the University Committee on Animal Resources (UCAR).

### Inhalation Exposure Paradigm

Whole-body inhalation exposures were conducted in the University of Rochester Inhalation Exposure Facility for 4hr/day from 8:00 A.M. 5 days a week for 12 weeks to mimic an agricultural work schedule during harvest season. During the exposure, mice were singly housed in wire-mesh chambers inside a 30-L steel-reinforced Lexan exposure chamber which was maintained at 40–45% relative humidity and 22–24°C in a fume hood. PQ aerosols were generated by adding a 0.2 mg/mL solution of PQ in ddH2O (made fresh daily) into an ultrasonic nebulizer (Ultrasonic2000, Nouvag Dental and Medical Equipment). Clean, dry air (approximately 500 mL/min) was passed through the nebulizer at a frequency of 2.4 MHz to produce a PQ mist, which then passed through a heated drying tube and cold trap to remove moisture. The subsequent PQ particles were mixed with diluting air and passed into the exposure chamber at a rate of 25–30 L/min. Using a peristaltic pump (Masterflex, Cole-Parmer Inc.), the PQ solution was slowly transferred into the nebulizer for the duration of the exposure (approximately 18 mL/hour). A condensation particle counter (CPC Model 3022A, TSI Inc.) was used to capture real-time exposure measurements, enabling quantification of continuous particle concentration. The particle size distribution of the aerosol was measured daily using an electrostatic classifier (SMPS Model 3071, TSI Inc.). Three filters (Pallflex Membrane Filters, Pall Life Science) were collected hourly, starting after the first 30 minutes of exposure and ending before the last 30 minutes. The gravimetric exposure concentration by mass was measured by weighing filters before and after use. The concentration of PQ on the filters was quantified chemically using a UV-Vis spectrophotometer (HP-Agilent Technologies) at an absorbance wavelength of 258 nm. Analytical standard grade PQ (PQ dichloride hydrate PESTANAL®, ≥98.0% purity) was purchased from Millipore Sigma (CAS No: 75365-73-0).

#### PQ quantification

PQ concentrations in the olfactory bulb, cortex, and brain stem were quantified by the University of Rochester Mass Spectrometry core to assess PQ uptake and distribution following inhalation. Frozen brain tissues were thawed, weighed, and diluted in 12% acetic acid spiked with 500 pg/mL of the internal standard, PQ D8 (Paraquat dichloride-[rings-d8] hydrate, PESTANAL® ≥98% purity, Millipore Sigma, Prod No: 50636), and homogenized for 10 seconds via ultra-sonication (SLPe digital sonifier, Branson Ultrasonics). The resulting homogenized solution was boiled for 10 minutes, then centrifuged at 10,700 rpm (12,000 g) at 4°C for 90 minutes, and the supernatant collected for analysis.

PQ analysis was performed using a Dionex Ultimate 3000 UHPLC system coupled to a Q Exactive Plus mass spectrometer (Thermo Fisher Scientific). PQ and its internal standard PQD8 were separated on an Agilent InfinityLab Poroshell 120 HILIC-Z column (2.1 × 100 mm). The mobile phase consisted of (A) 20 mM ammonium formate in 0.1% formic acid and (B) 20 mM ammonium formate in 90% acetonitrile with 0.1% formic acid. The flow rate was maintained at 800 µl/min, and the column temperature set to 30°C. A 10 µl injection volume was used for each sample. Analytes were eluted using a 4-minute gradient starting at 80% mobile phase A, held for 0.5 minutes, followed by a ramp to 60% mobile phase B over 0.5 minutes. This composition was held for 0.5 minutes, then returned to 80% B over 0.25 minutes, with re-equilibration for an additional 2.25 minutes.

The Q Exactive Plus operated in positive ion mode using a heated electrospray ionization (HESI) source. Source settings included a spray voltage of 4.0 kV, a sheath gas flow rate of 50, an auxiliary gas flow rate of 10, and a capillary temperature of 350°C. Detection was performed using a parallel reaction monitoring (PRM) method. Precursor ions of 186.1153 m/z for PQ and 194.1645 m/z for PQD8 were isolated with a 1.0 m/z window and subjected to higher-energy collisional dissociation (HCD) with a collision energy of 25. The maximum injection time was set to 200 ms, and the automatic gain control (AGC) target was 2 × 10⁵. Fragment ions were acquired in the Orbitrap at a resolution of 17,500 (at m/z 200). Quantification was based on fragment ions 171.0916 m/z for PQ and 179.1408 m/z for PQD8. Chromatographic peaks were extracted with a 10 ppm mass tolerance and integrated using the LC Quan node in XCalibur software (Thermo Fisher). Final concentrations were calculated by comparing PQ:PQD8 peak area ratios against a standard curve.

### Locomotor Behavioral Assays

#### Open Field Test

Spontaneous movement was measured using a VersaMax (monitor RXYZCM-16, analyzer, VMAUSB; Accuscan, Columbus, OH). The mice were given a 2-minute acclimatization period followed by a 60-minute test session. This assay measured parameters of ambulation and stereotypic behavior at the edges and the center of the open field.

#### Grip Strength

A digital force gauge (Columbus Instruments, Columbus, OH) was used to assess peak force applied on a digital display with a grid that allowed the mouse to grip by either or both paws. Each mouse was lifted by the tail to a height where the front paws were at the same height as the bar and then moved horizontally toward the bar until it was within reach. Once the grip was deemed symmetric, the mouse was pulled away at a constant and slow speed until its grasp was broken. Measurements were discarded if the animal used only one paw or also used its hind paws, turned backwards during the pull, or left the bar without resistance. Each mouse was tested three times on both forelimbs and hindlimbs, with a minimum of 60 seconds between trials.

#### Inverted Screen

Mice were placed on a 4 x 4-inch screen of #4 mesh supported by a frame and attached to a metal rod. Once the mouse firmly grasped the mesh, it was rotated 180 degrees such that the mouse was turned upside down. Time required for the mice to right itself by climbing up the mesh was measured. Each mouse was tested three times, non-consecutively.

### Neurotransmitter Analysis

Neurotransmitter concentrations were quantified in the University of Rochester Mass Spectrometry Core using micro-dissected striata from the non-behavior cohort. The targeted analytes encompassed dopaminergic (tyrosine [Tyr], dopamine [DA], 3,4-dihydroxyphenylacetic acid [DOPAC], homovanillic acid [HVA], norepinephrine [NE]), serotonergic (tryptophan [Trp], kynurenine [Kyn], 5-hydroxytryptophan [5-HTP], serotonin [5-HT], 5-hydroxyindoleacetic acid [5-HIAA]), and glutamatergic (glutamine [Gln], glutamate [Glu], γ-aminobutyric acid [GABA]) neurotransmitters. Amino acids not typically considered classical neurotransmitters involved in energy metabolism and protein turnover were also measured (alanine [Ala], valine [Val], leucine[Leu], isoleucine [Ile], phenylalanine [Phe], arginine [Arg], histidine [His], serine [Ser], threonine [Thr], asparagine [Asn], proline [Pro], lysine [Lys]).

Tissue samples were thawed from −80°C storage, weighed, and homogenized in 75 μL of ice-cold 50% acetonitrile (v/v) using a Branson SLPe digital sonifier (Branson Ultrasonics Corp., Danbury, CT, USA) with 10 ultrasound pulses. Homogenates were incubated on ice for 10 minutes, followed by centrifugation at 10,000 × g for 20 minutes at 4°C. The resulting supernatants were collected and subjected to a second centrifugation under identical conditions. Final supernatants were transferred into clean tubes and stored at −80°C until further analysis.

Stock solutions of all neurotransmitters (Sigma-Aldrich, St. Louis, MO, USA) were prepared at 5 mg/mL. Tyr was dissolved in 0.2 M HCl, while all other neurotransmitters were dissolved in deionized water. Analyte concentrations in the standard mix were selected based on prior range-finding experiments and known regional differences in endogenous neurotransmitter levels. Internal standards were generated following a modified derivatization method adapted from another paper [86] using 13C₆-benzoyl chloride (BzCl; Sigma-Aldrich) as the derivatizing agent. Once derivatized, internal standard mixtures were aliquoted and stored at −80°C for long-term use.

Internal standard aliquots were thawed and diluted for sample preparation in 50% acetonitrile containing 1% sulfuric acid. Samples were then derivatized using the same procedure: after centrifugation at 16,000 × g for 5 minutes to remove debris, 20 μL of supernatant was transferred into LoBind Eppendorf tubes, followed by sequential addition of 10 μL of 100 mM sodium carbonate, 10 μL of 2% BzCl in acetonitrile, and 10 μL of internal standard. To reduce organic content before LC-MS injection, 50 μL of deionized water was added. A final centrifugation was performed to pellet residual protein, and the supernatant was transferred to clean autosampler vials.

Liquid chromatography–mass spectrometry (LC-MS/MS) analysis was performed using a Dionex Ultimate 3000 UHPLC system coupled to a Q Exactive Plus mass spectrometer (Thermo Fisher Scientific, Waltham, MA, USA). Separation was achieved on a Waters Acquity HSS T3 column, with mobile phases consisting of (A) 10 mM ammonium formate in 0.1% formic acid and (B) acetonitrile. The flow rate was 400 μL/min, and the column oven temperature was maintained at 27°C. Following the injection of 5 μL per sample, analytes were eluted using a 12-minute multi-step gradient. The mass spectrometer operated in positive ion mode. Derivatized analytes were detected using parallel reaction monitoring (PRM), and fragment ions were extracted with a 10 ppm mass accuracy window using the LC Quan module within XCalibur software (Thermo Fisher Scientific). Relative quantification was performed by comparing the peak areas of endogenous analytes to their respective internal standards. Neurotransmitter concentrations in each sample were normalized to the wet tissue weight and expressed as a percentage of the air-exposed control group by dividing each PQ-treated sample value by the control group mean for that analyte.

### Western Blot

Tissue lysates were prepared using radioimmunoprecipitation assay (RIPA) buffer (Thermo Scientific). The supernatants were obtained after centrifuging the homogenates at 15,000 rpm for 1 hour. Protein concentration was measured using Bradford dye (Bio-Rad). Lysates containing equal amounts of protein were separated on a 15% weight/volume SDS-polyacrylamide gel (Biorad, Cat No. 4561084). After separation, proteins were electroblotted onto a nitrocellulose membrane, and nonspecific-binding sites were blocked with a 10% BSA blocking buffer. TH (1:1000; EMD Millipore, Cat. AB152) and GAPDH (1:1000; ORIGENE, Cat. TA337137) primary antibodies were used to blot the membranes overnight (16 hours) at 4°C. Blots were washed in PBS containing 0.01% Tween-20 and incubated for 1 hour in the respective secondary antibodies.

### Immunofluorescent Staining

Immunofluorescent staining was performed as described previously [87] with brief modifications. Brains were sectioned to a thickness of 50 μm on the coronal plane via freezing sliding microtome and stored in cryoprotectant at −20C until ready for use. Tissue sections were mounted onto slides, antigen retrieved in 1X citrate buffer pH 6.0 (Sigma-Aldrich; catalog no. C9999) at 99C for 30 minutes, cooled for 1 hour at room temperature, permeabilized with 0.5% Triton X-100 in 1X PBS for 30 minutes, blocked with 10% BSA/0.1% Triton X-100 in 1X PBS for 1 hour at room temperature, and incubated overnight at 4C with Rabbit-anti-Alpha-synuclein [MJFR1] 1:100 (Abcam; catalog no. 138501) and Sheep-anti-tyrosine hydroxylase 1:1000 (Millipore Sigma; catalog no. AB1542) primary antibodies diluted in 1% BSA/0.1% Triton-X 100 in 1X PBS. The following day, the tissue was washed and incubated with Donkey-anti-rabbit Alexa Fluor Plus 488 (Invitrogen; catalog no. A3279) and Donkey-anti-sheep Alexa Fluor 594 (Invitrogen; catalog no. A11016) secondary antibodies diluted at 1:1000 in in 1% BSA/0.1% Triton-X 100 in 1X PBS for 2 hours at room temperature, followed by TrueBlack Lipofuscin Autofluorescence Quencher (Biotium; catalog no. 23007) for 10 minutes at room temperature, then cover slipped with ProLong Gold antifade reagent with DAPI (Invitrogen; catalog no. P36935). Tissue was imaged at 60X magnification using Keyence BZ-X810 microscope (Keyence; Osaka, Japan), and fluorescence intensity was analyzed using the Keyence BZ-X800 Analyzer software (Keyence; Osaka, Japan).

### Immunofluorescence Analysis

#### Substantia nigra

60X magnification z-stacked images were acquired using the Keyence BZ-X810 microscope. 5-8 images across 3-4 tissue sections (Bregma −2.9mm to −3.5mm), 200 μm apart, were taken per sample. Z-stacked images were compressed into a single full-focused image, and total alpha-synuclein fluorescence and area intensity were measured in TH-positive cells and DAPI-stained nuclei of the substantia nigra using the single-extraction colocalization plugin on the Keyence BZ-X800 Analyzer software. Total TH fluorescence and area intensity per cell were also measured for each section using the Keyence BZ-X800 Analyzer software. Data are represented as brightness or area per cell.

#### Olfactory bulb

60X magnification z-stacked images were acquired using the Keyence BZ-X810 microscope. 8-12 images across 3-4 tissue sections (Bregma +4.3mm to +3.7mm), 200 μm apart, were taken per sample. Z-stacked images were compressed into a single full-focused image. Total alpha-synuclein and TH fluorescence and area intensities were measured in the glomerular layer of the olfactory bulb using the Keyence BZ-X800 Analyzer software. Total synuclein and TH data are represented as total brightness or area per section. Nuclear alpha-synuclein fluorescence and area intensity were measured in DAPI-stained nuclei using the single-extraction colocalization plugin on the Keyence BZ-X800 Analyzer software. Nuclear alpha-synuclein data are represented as brightness or area per DAPI count.

### Immunohistochemistry and Unbiased Stereology

Immunohistochemistry and unbiased stereology were performed essentially as described previously [88]. Briefly, the midbrain was coronally sectioned (50 μm) with a freezing sliding microtome and stored in cryoprotectant at −20 °C until ready for use. Free-floating midbrain sections (every 4^th^ section from Bregma −2.60 to −3.80) encompassing the Substantia Nigra *pars compacta* (SNc) were stained for TH. Sections were treated with 75% methanol/2.5% H_2_O_2_ in water for 20 minutes to quench endogenous peroxidase, permeabilized with 1% Triton X-100 in 1X PBS for 15 minutes, blocked with 10% BSA/0.1% TX100 for 1 hour at room temperature, and incubated with Rabbit-anti-TH (Millipore Sigma; catalog no. AB152) at a dilution of 1:500 in 1% BSA/0.1%TX100 in 1X PBS for 48 hours at 4°C. The sections were then washed with 1X PBS and incubated with biotinylated Goat-anti-Rabbit IgG (Vector Laboratories; catalog no. BA-1000) secondary antibody at a dilution of 1:200 in 1% BSA/0.1% Triton X-100 in 1X PBS for 2 hours at room temperature, followed by incubation of the avidin-peroxidase complex (Vector Laboratories catalog no. PK-4000) for 1 hour at room temperature, then developed using DAB substrate (Vector Laboratories; catalog no. SK-4100).

Unbiased stereology counting was performed by a blinded individual using the Leica DM4 B LED microscope and Microbrightfield Stereo Investigator (MBF Biosciences) computer-assisted stereology software. Unilateral counts of TH^+^ cells were performed on every 4^th^ section, for a total of 4-5 sections per sample, throughout the SNc. The SNc borders were determined using a 5X objective lens, and were counted at 40X magnification using the optical fractionator probe. A counting frame of 100 μm x 100 μm with a framing grid size of 250 μm x 250 μm and a disector height of 20 μm was chosen. Only cells that came into focus within the counting frame were counted. The coefficient of error (Gundersen, m=1) for all samples was equal to or below 0.1, and 100-200 markers were counted within a minimum of 40 framing sites for each sample.

### Statistical Analysis

Given that previous studies have demonstrated sex specific differences in the behavioral and biochemical effects in PQ models [17, 89, 90], statistical analyses were separated by sex. Data were analyzed either using 2-factor ANOVA, with time and PQ as variables, or using t-tests to determine differences in treatment. All statistical analyses were performed using JMP (JMP Statistical Discovery, Inc.), and p-values ≤ 0.05 were considered statistically significant.

